# hu.MAP 2.0: Integration of over 15,000 proteomic experiments builds a global compendium of human multiprotein assemblies

**DOI:** 10.1101/2020.09.15.298216

**Authors:** Kevin Drew, John B. Wallingford, Edward M. Marcotte

**Author notes:** To whom correspondence should be addressed: KD, EMM.

## Abstract

A general principle of biology is the self-assembly of proteins into functional complexes. Characterizing their composition is, therefore, required for our understanding of cellular functions. Unfortunately, we lack a comprehensive set of protein complexes for human cells. To address this gap, we developed a machine learning framework to identify protein complexes in over 15,000 mass spectrometry experiments which resulted in the identification of nearly 7,000 physical assemblies. We show our resource, hu.MAP 2.0, is more accurate and comprehensive than previous resources and gives rise to many new hypotheses, including for 274 completely uncharacterized proteins. Further, we identify 259 promiscuous proteins that participate in multiple complexes pointing to possible moonlighting roles. We have made hu.MAP 2.0 easily searchable in a web interface (http://humap2.proteincomplexes.org/), which will be a valuable resource for researchers across a broad range of interests including systems biology, structural biology, and molecular explanations of disease.

## Introduction

Macromolecular protein complexes carry out a wide variety of functions in the cell including essential functions such as replication, transcription, translation and protein degradation (e.g. MCM-ORC, RNA polymerase, ribosome, proteasome)^1,2^. The disruption of protein complexes is implicated in many human diseases^3,4^ and many therapeutics target protein complexes^5^. The formation of protein complexes to carry out biological function is a general principle of biology and characterizing their composition is therefore a basic requirement to our full understanding of cellular functions. Unfortunately, we still lack a comprehensive set of protein complexes for the human cell.

To address this gap in knowledge, there is an ongoing worldwide effort to identify all protein interactions in human cells. High throughput protein interaction screens using affinity purification (AP-MS)^6–9^ have greatly increased coverage of protein interactions across the proteome. Likewise, biochemical separation strategies, such as co-fractionation mass spectrometry (CF-MS), have provided orthogonal approaches to identifying protein complexes^10–13^. Although these methods have identified tens of thousands of protein interactions, they still have limited coverage of the entire human interactome.

Fortunately, these high throughput methods are orthogonal, each sampling different parts of the human proteome and identifying nonoverlapping sets of interactions. We previously re-analyzed and integrated three of the largest datasets available at the time^6,7,10^, over 9,000 mass spectrometry experiments, to build a more complete and accurate set of protein complexes^14^. Our resource, hu.MAP, identified interactions for over a third of all human proteins.

We envision interactomes as evolving entities, growing and improving as new technologies and datasets emerge. Here, we introduce hu.MAP 2.0, which we find to be the most accurate and comprehensive human protein complex map available to date. hu.MAP 2.0 is an integration of over 15,000 mass spectrometry experiments and identifies 6,969 complexes consisting of 57,178 unique interactions among 9,968 human proteins. Multiple performance metrics demonstrate that hu.MAP 2.0 outperforms our previous map as well as several other complex maps available in the literature. We further show that complexes in hu.MAP 2.0 are not only highly enriched for specific literature-curated annotations but also have greater coverage of completely uncharacterized genes. Finally, we highlight several new biological findings that illustrate the utility of hu.MAP 2.0 as a resource for biological discovery.

## Methods

Our strategy in building a comprehensive map of protein complexes involves the integration of the many orthogonal experimental protein interaction datasets available using a custom machine learning pipeline as shown in **Figure 1A**. Each individual experimental dataset identifies nonoverlapping sets of protein interactions and therefore combining them results in a more accurate and comprehensive set of interactions. Our pipeline combines quantified features from these datasets using an SVM classifier which calculates a confidence score of two proteins interacting. This results in a large protein interaction network. The network is subsequently searched for dense regions of highly connected proteins which represent individual complexes. The identified complexes are ranked by a clustering confidence value and we call the resulting set of complexes, hu.MAP 2.0. The map of complexes contains many known complexes such as EIF2B complex, Spliceosome, RNA Pol III and IFT-A complex (**Figure 1A** Positive Control Examples) as well as many novel insights into the physical biology of the cell.

**Figure 1.**
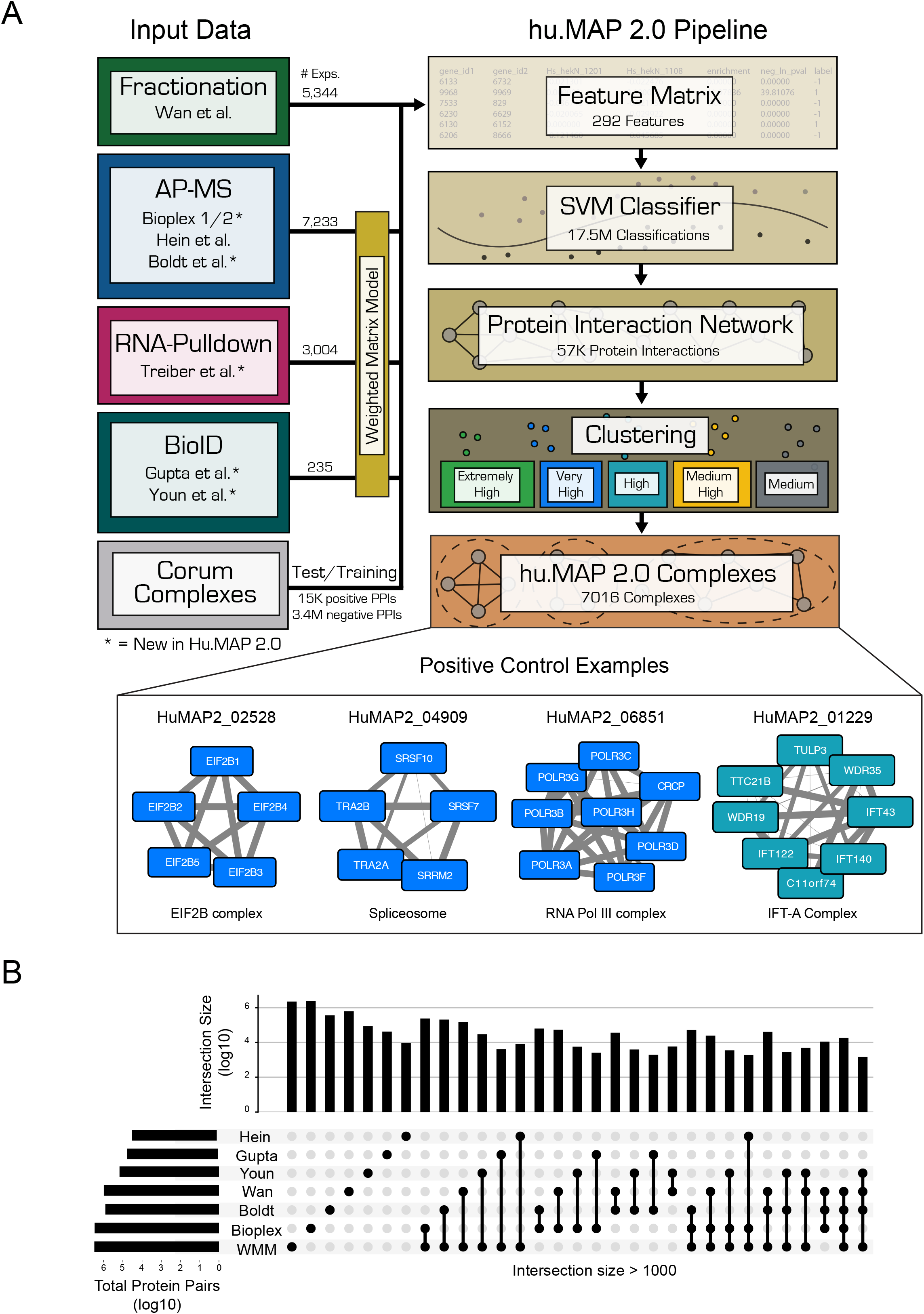
Machine learning framework to identify protein complexes. **(A)** Graphical description of computational pipeline to integrate >15,000 mass spectrometry experiments. A Support Vector Machine (SVM) classifier was trained using numerical measures (i.e. features) on pairs of proteins calculated from original mass spectrometry data and training labels from literature curated complexes (CORUM). The classifier was then used to construct a protein interaction network by calculating a confidence score for all pairs of proteins for their propensity to interact. Clustering parameters were then learned from training complexes and five final clusterings were chosen ranked in order of confidence from “Extremely High” to “Medium”. The union of these selected clusterings represent the final set of hu.MAP 2.0 complexes. Networks of previously known protein complexes identified by this pipeline which were not in the training set of complexes are shown as positive control examples. **(B)** “UpSet” plot displaying the intersections of protein pairs for all integrated datasets. The plot shows the Weighted Matrix Model (WMM) provides additional information for many pairs of proteins that would be limited otherwise.

### Integration of over 15,000 mass spectrometry experiments

To construct hu.MAP 2.0 we integrated over 15,000 previously published mass spectrometry experiments using our custom machine learning framework. We built upon the 9,000 mass spectrometry experiments used for hu.MAP 1.0 ^6,7,10,14^ by incorporating additional affinity purification data from Bioplex 2^8^ and Boldt et al.^15^ as well as proximity labeling data from Gupta et al.^16^ and Youn et al.^17^ (**Figure 1A**). The “Upset” plot in **Figure 1B** shows that tens of thousands of protein pairs are represented by 2 or more datasets providing orthogonal evidence for those interactions. This greatly enhances our frameworks ability to identify true interactions from false ones.

Additionally, we applied our Weighted Matrix Model (WMM) technique which we previously demonstrated identifies many new high confidence interactions from affinity purification data^14,18^. Contrary to the traditional spoke and matrix models used to interpret AP-MS data that only consider one purification experiment at a time, the WMM technique takes all experiments into account and determines protein pairs that are in the same experiments more often than random chance. The WMM balances both the false negative and false positive issues that face both the spoke and matrix models and therefore is capable of identifying novel interactions. Also in contrast to the traditional models, the WMM can be applied to datasets that were not exclusively collected for the purpose of identifying protein interactions. Specifically, we applied our WMM to >3,000 RNA hairpin pulldown experiments^19^ and incorporated the results into our framework. **Figure 1B** shows WMM overlaps substantially with other methods but also provides evidence of interactions between many pairs of proteins not covered in the other datasets.

### hu.MAP 2.0 network is highly accurate

We next trained an SVM classifier to determine whether two proteins interact in a macromolecular complex. The classifier uses 292 features computed from the mass spectrometry experiments and is trained on examples consisting of co-complex protein interactions from a set of >1,100 literature curated Corum complexes^20^. The set of Corum complexes were split into two equal proportion non-overlapping test and train subsets. We then derived sets of co-complex interactions from these test and train complex subsets. The classifier was then parameterized using 5x cross-validation on the train co-complex interactions and applied to 17.5 million pairs of proteins scoring each one for their ability to interact based on the input data (see **Supplemental Methods**). The full list of scored pairs can be found here: http://humap2.proteincomplexes.org/download.

To evaluate the performance of our SVM classifier, we examined the confidence scores of 8,337 co-complex interactions in our leave-out test set using a Precision-Recall framework (**Figure 2A**). Precision-Recall frameworks are the preferred method of evaluating imbalanced binary classification problems such as protein interaction identification^21,22^. We see that hu.MAP 2.0 outperforms our previous complex map, hu.MAP 1.0, as well as all other datasets including Bioplex2, Wan et al. and Hein et al., demonstrating the power of data integration. Additionally, we see a drop in performance when we remove the WMM features from hu.MAP 2.0 further showing how important these features are to performance improvements. Finally, we compare hu.MAP 2.0 to the HuRi dataset which encompasses a 17,500 × 17,500 all by all yeast2hybrid screen^23^ and show the HuRi network, which aims to capture only direct protein-protein interactions, underperforms all other networks when evaluated on co-complex interactions.

**Figure 2.**
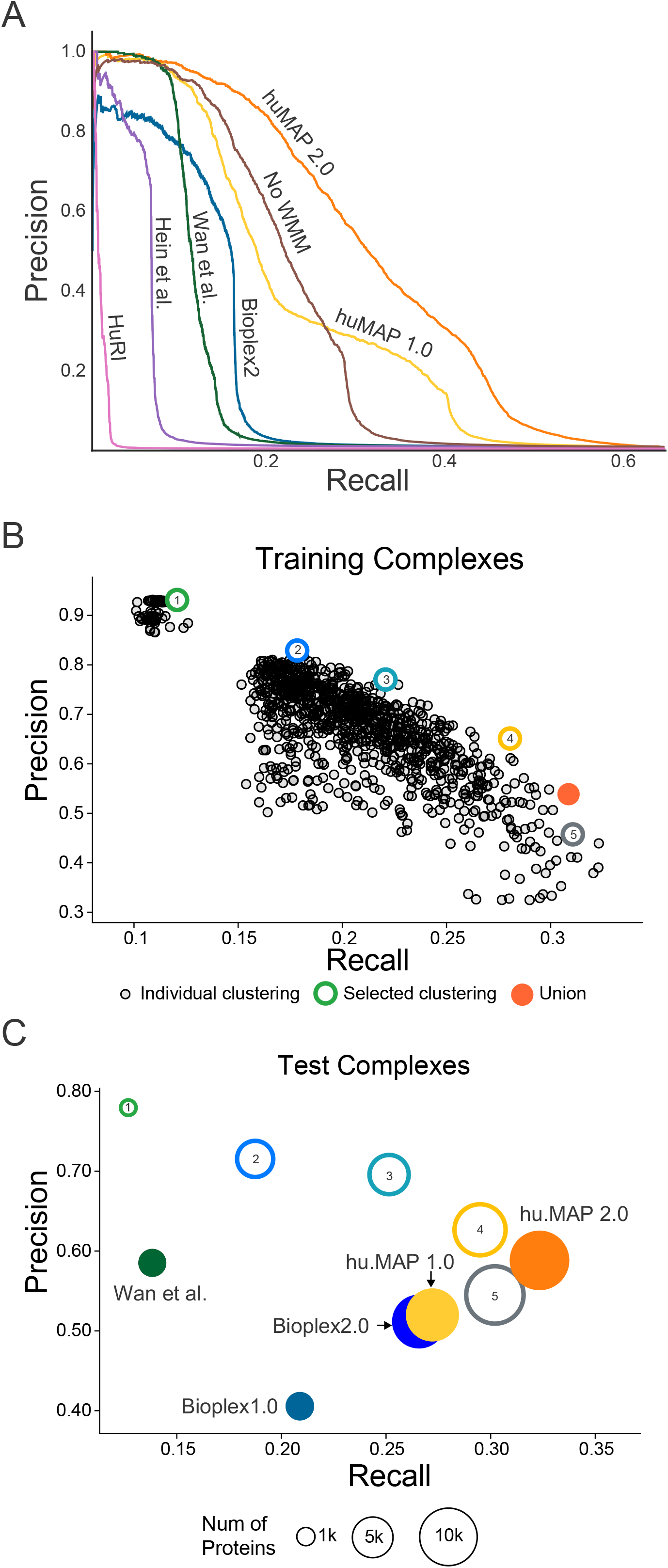
hu.MAP 2.0 outperforms previous complex maps. **(A)** Precision-Recall (PR) plot evaluated on a test (i.e. leave-out) set of literature curated co-complex pairwise protein interactions. The plot shows hu.MAP 2.0 is more accurate and comprehensive than previous published datasets. The plot also evaluates the performance of predictions without the Weighted Matrix Model (WMM) and shows the WMM substantially improves performance. **(B)** PR scatter plot of 1,700 clustering parameter sets. Individual clusterings were evaluated using the *k*-clique method (see **Supplemental Methods**) on training complexes. Five clusterings (colored hollow circles) were selected representing varying degrees of confidence balancing the tradeoff between precision and recall. The five selected clusterings were combined as a final set of clusters (orange filled circle). **(C)** PR scatter plot of hu.MAP 2.0 complexes (orange filled circle) and other published complex maps (colored filled circles) evaluated on a test set of literature curated complexes. hu.MAP 2.0 complexes increase in both precision and recall relative to other maps. Also plotted are the five sets of complexes at different levels of confidence (colored hollow circles) demonstrating consistency between the level of confidence determined from training set **(B)** and test set.

### hu.MAP 2.0 complex map

Once we confirmed our protein interaction network was of high quality, we then clustered this network using a parameterized two-stage clustering algorithm to identify highly connected proteins which represent protein complexes. Briefly, the algorithm first uses the ClusterOne algorithm^24^ to cluster the entire network and then each resulting cluster is further clustered using the MCL algorithm^25^. There are several parameters for each algorithm that require optimization in addition to a parameter representing the confidence threshold of the input network. We therefore optimize these parameters by generating clusterings for over 1,700 parameter combinations and evaluated each clustering’s performance using the *k*-clique precision-recall performance measure^14^ (**Figure 2B**). The resulting clusterings vary substantially with regards to their performance but ultimately show a familiar pattern of a tradeoff between precision and recall. We therefore selected 5 clusterings that balance this tradeoff. For example, clustering 1 (green) is a set of clusters with “extremely high” precision but low recall (1) while clustering 5 (grey) is a set of clusters with “medium” precision yet higher recall (**Figure 1A**, **Figure 2B**). We then combined all 5 selected clusterings into a union set while preserving their precision rank (**Supplemental Table 1**).

**Table 1:**
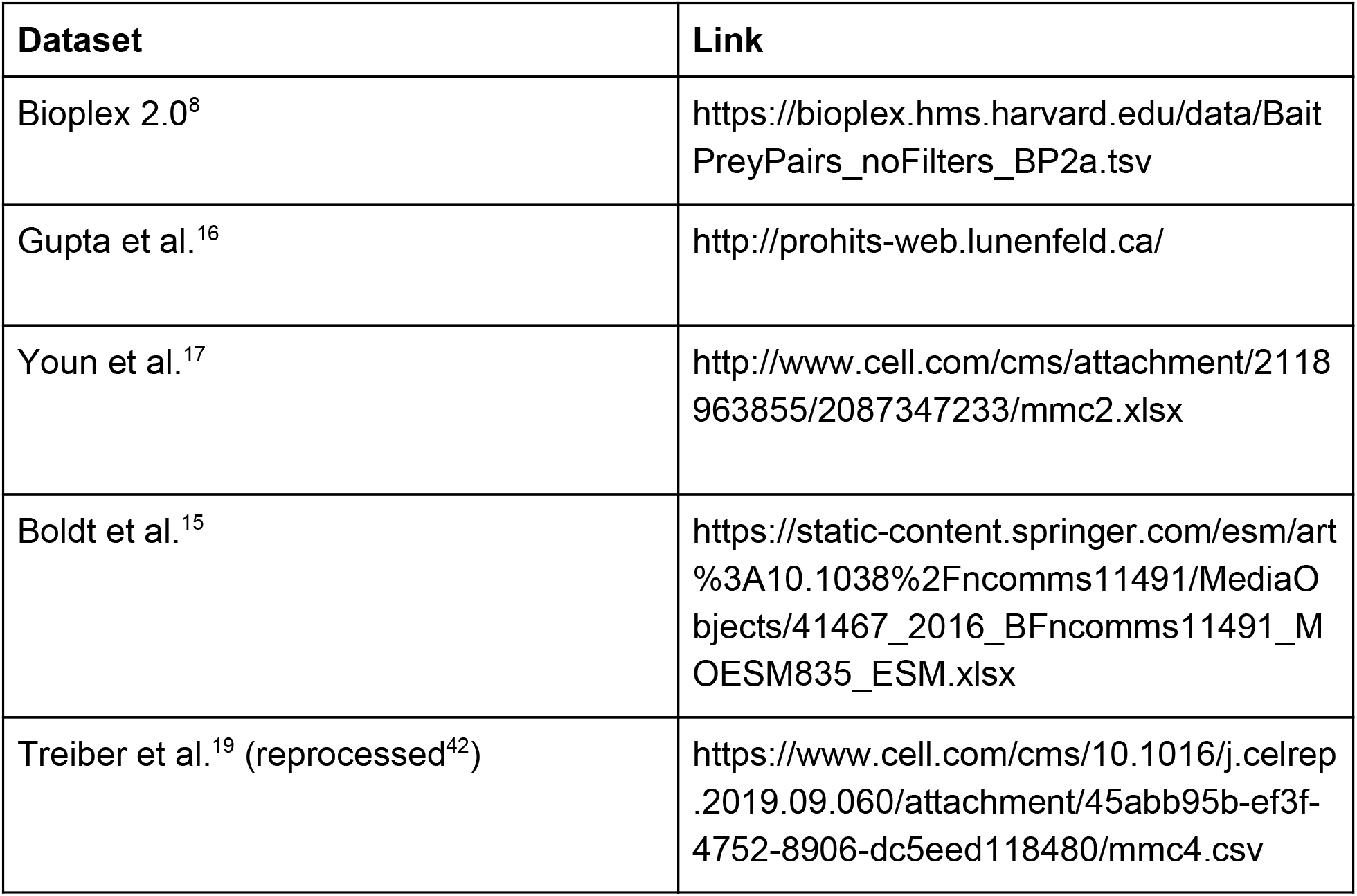
Dataset Raw Features

To evaluate the union of clusterings that represent our final hu.MAP complexes, we again use the *k*-clique precision recall performance measure but now calculated on the leave-out set of test complexes. As shown in **Figure 2C**, our hu.MAP 2.0 complexes balances both precision and recall. Additionally, we see a consistent trend of *k*-clique precision and recall values for our individual clusterings between both the train (**Figure 2B**) and test sets (**Figure 2C**). This suggests the confidence ranking given to each complex is robust. We also observe our final set of complexes outperforms our previous hu.MAP 1.0 complexes as well as other previous state-of-the-art complex maps. Taken together, this points to hu.MAP 2.0 complexes as being highly accurate and spanning a large portion of all human protein assemblies.

## Results and Discussion

### Identification of multifunctional promiscuous proteins

Now that we have established hu.MAP 2.0 as a highly accurate resource of human protein complexes, we can ask questions that were previously hindered by less accurate maps. Specifically, one question that has eluded researchers is: how prevalent are promiscuous proteins in stable protein assemblies? That is, how often do we see proteins participating in multiple different complexes (sometimes termed “moonlighting”) and presumably performing orthogonal functions. We therefore created a non-redundant set of complexes (see **Supplemental Methods**), and surprisingly identified 259 proteins that participate in multiple complexes (**Supplemental Table 2**). This constituted nearly 7.5% of proteins in the non-redundant set of complexes.

**Table 2:** Selected Clusterings Parameters

| Clustering | Confidence | Score Threshold | ClusterOne Density | ClusterOne Overlap | MCL Inflation |
| --- | --- | --- | --- | --- | --- |
| 1 | Extremely High | 1.0 | 0.4 | 0.6 | 9 |
| 2 | Very High | 0.7 | 0.4 | 0.6 | 9 |
| 3 | High | 0.5 | 0.4 | 0.7 | 4 |
| 4 | Medium High | 0.04 | 0.4 | 0.7 | 2 |
| 5 | Medium | 0.02 | 0.1 | 0.6 | 2 |

**Figure 3A** shows an example of a promiscuous protein, HSPA9, participating in two unrelated protein complexes. HSPA9 is a multilocational (mitochondria and nucleus) and multifunctional protein, playing a role in mitochondrial import as well as stress response^26^. We identify HSPA9 participating in two complexes that reflect its multifunctional role, specifically HuMAP2_01130, a heat shock response complex and HuMAP2_00358, a mitochondrial protein import complex. To further verify HSPA9’s membership in these two complexes, we inspected sparkline traces of two co-fractionation experiments^10^ (**Figure 3A, bottom**). We identified two separate elution peaks of HSPA9 which correspond to the two complexes. This example demonstrates the ability of our complex map to identify multifunctional promiscuous proteins and place them into their respective non-overlapping functional complexes.

**Figure 3.**
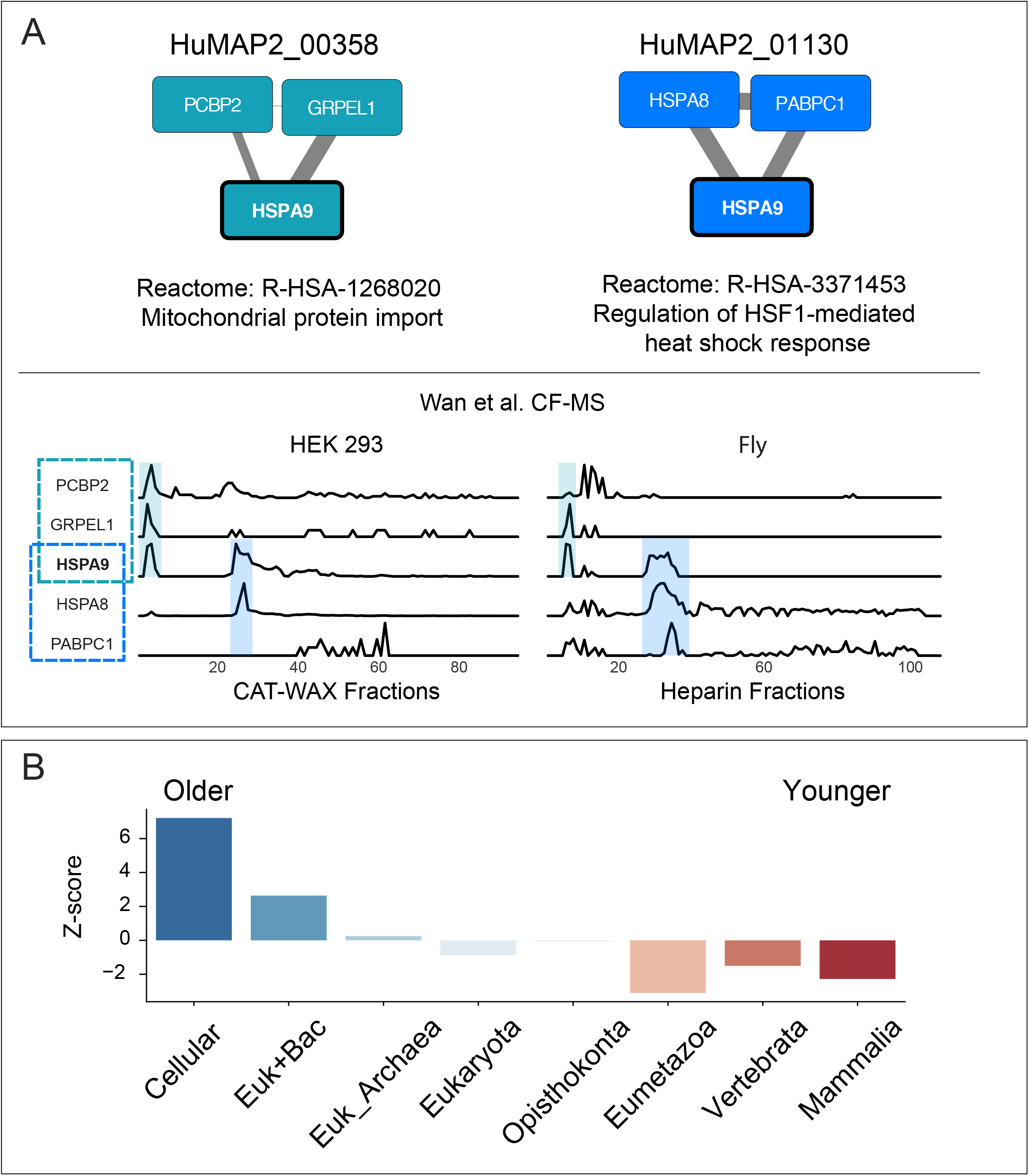
hu.MAP 2.0 complexes identify promiscuous proteins. **(A, top)** Multifunctional protein HSPA9 participates in two distinct complexes, HuMAP2_00358 and HuMAP2_01130. HuMAP2_00358 (turquoise, “high” confidence) is enriched for Reactome annotation “Mitochondrial protein import”, a known function of HSPA9. HuMAP2_01130 (blue, “very high” confidence) is enriched for Reactome annotation “Regulation of HSF1-mediated heat shock response”, another known function of HSPA9. Weight of network edges represent confidence of interactions. **(A, bottom)** Sparkline elution profiles from two orthogonal biochemical fractionation experiments. HEK 293 cell lysate was separated using a mixed bed ion-exchange column and D. melanogaster embryo lysate was separated using a heparin column^10^. HSPA9 elutes in two distinct peaks which co-elute with members of the two complexes. **(B)** Promiscuous proteins are older on average than single complex proteins. Z-scores for each age group were determined by comparing the number of promiscuous proteins to a randomly sampled background set consisting of non-promiscuous proteins (i.e. participating in only 1 complex).

We next hypothesize that these promiscuous proteins would be on average older due to younger proteins not having enough evolutionary time to make multiple connections. **Figure 3B** shows a clear enrichment in older proteins in the set of promiscuous proteins as well as a depletion for younger proteins (see Supplemental Methods). Using gProfiler^27^ to identify functionally annotations for the older promiscuous proteins, we identify older promiscuous proteins are enriched for Reactome “Metabolism” (adjusted p-value ~5×10^-8) among other metabolism related annotations (**Supplemental Figure 1**). Consistent with this finding, many “moonlighting” proteins are enzymes with multifunctional roles^28^. Our results suggest a protein’s complex membership may play a role in its multifunctional activity.

### Functional annotation of uncharacterized proteins

High quality protein complex maps have long been sought after for the purpose of functionally annotating poorly characterized proteins in a genome^29–31^ due to the relationship between physical interaction and biological function. To assess hu.MAP 2.0’s ability to be used to functionally annotate uncharacterized proteins, we first tested whether our identified complexes are enriched with literature-curated annotations including Gene Ontology^32^, Reactome^33^, Corum^20^, Human Phenotype Ontology^34^, and KEGG^35^. We see in **Supplemental Figure 2** that >40% of our complexes are enriched with at least one annotation which is 20.5 fold higher than expected by randomly shuffled complexes. This result shows hu.MAP 2.0 complexes are functionally coherent.

As an example of the utility of hu.MAP 2.0 in annotating poorly characterized proteins, **Figure 4A** shows two previously unreported interactions with RNaseH2, CMTR1 and SETD3. The evidence for these interactions are elucidated from co-fractionation mass spectrometry experiments which demonstrate a high degree of correlation over multiple separation column types and multiple organisms, which we have shown suggests a deep conservation in function^10^. RNaseH2 is implicated in Aicardi–Goutières syndrome, a monogenic autoinflammatory disorder which mimics *in utero* viral infection of the brain^36^. Mechanistically, RNaseH2 degrades RNA fragments of RNA-DNA hybrids including Okazaki fragment RNA primers during DNA replication^37^. Mutations in RNaseH2 are thought to disrupt the degradation of immuno-stimulating nucleic acids and cause innate immune activation^36^. CMTR1 is a mRNA methyltransferase and a known regulator of protein expression of IFN-stimulated genes to restrict viral infection^38^. SETD3, also a methyltransferase, is a human host protein critical for infection of a wide range of viruses and participates in viral replication yet its mode of action is currently unknown^39^. The links we identify between CMTR1, SETD3, and RNaseH2 point to potential mechanisms in which CMTR1 and SETD3 interact with RNaseH2 to modulate the innate immune response and affect viral replication.

**Figure 4.**
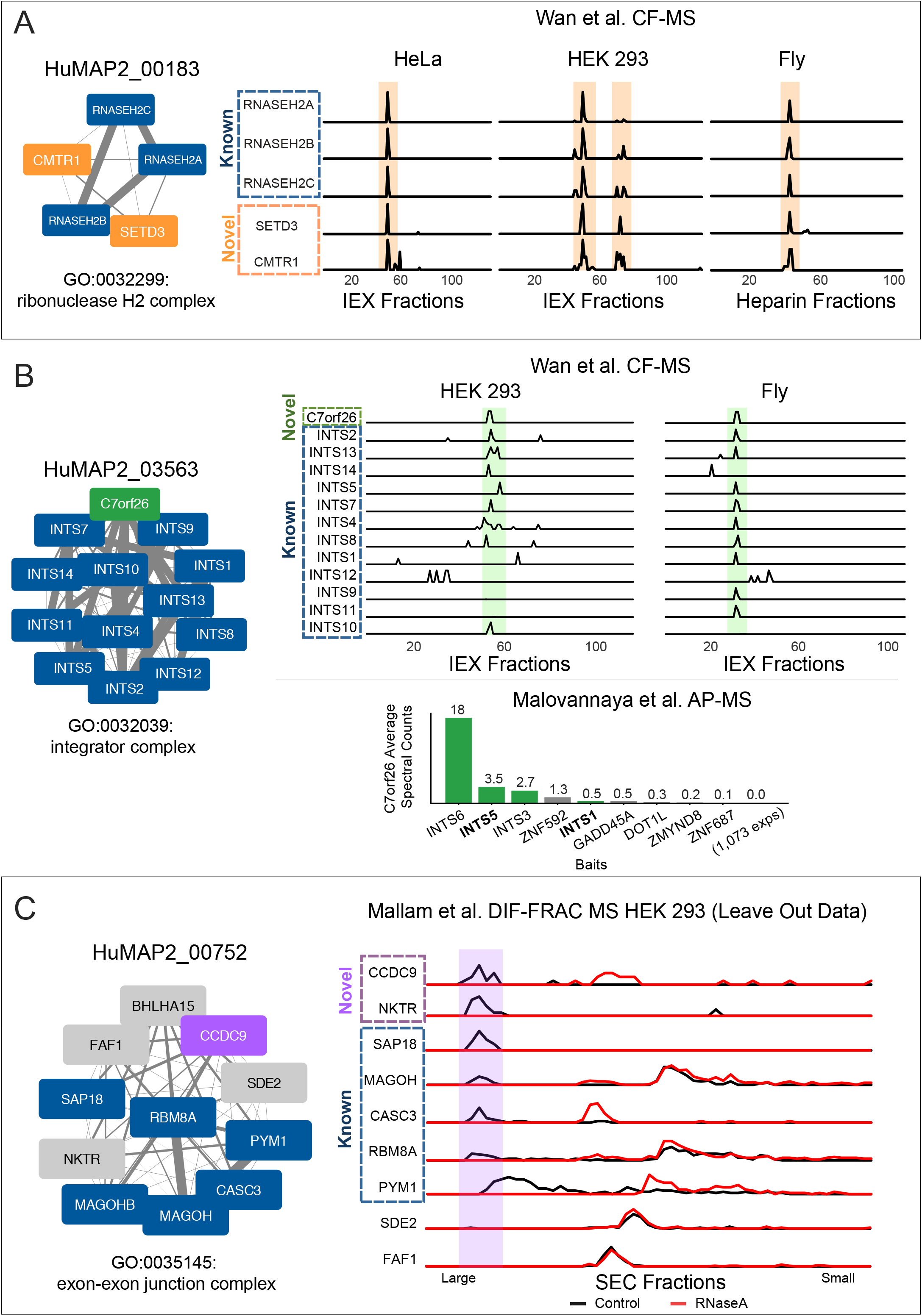
Transfer of function annotations to uncharacterized proteins. **(A)** SETD3 and CMTR1 are identified as co-complex interactors with the Ribonuclease H2 complex which provides a possible mechanistic explanation for their role in viral infection. Sparkline elution profiles from multiple orthogonal co-fractionation experiments demonstrate a strong degree of co-elution among subunits in the complex. Weight of network edges represent confidence of interactions. **(B)** The uncharacterized protein, C7orf26, is identified as part of the Integrator complex. Sparkline elution profiles show a high degree of correlation between C7orf26 and subunits of the Integrator complex from multiple orthogonal co-fractionation experiments. The association is additionally supported by affinity purification mass spectrometry (AP-MS) experiment where C7orf26 is pulled down with Integrator subunit baits. **(C)** The uncharacterized protein, CCDC9, is identified as co-complex with the exon-exon junction complex (EJC), a ribonucleoprotein complex involved in splicing. Sparkline elution profiles from the independently collected RNA DIF-FRAC size exclusion chromatography (SEC) experiment shows CCDC9 co-elutes with known subunits of the EJC when RNA is present (black). The elution profiles also show CCDC9 is sensitive to RNAse A treatment (shift of elution peak between black and red profiles) as are the subunits of the EJC further supporting CCDC9’s participation in this known ribonucleoprotein complex.

It has been observed that biomedical research is biased towards the study of well annotated genes and this bias is due less to the physiological importance or disease relevance of the gene but rather to the ease of experimentation using traditional methods^40^. Unbiased systematic approaches such as the integration of thousands of mass spectrometry experiments described here provide a powerful tool for closing the gap of uncharacterized proteins. We therefore cross-referenced genes deemed poorly annotated (Uniprot^41^ annotation score <= 3) with hu.MAP 2.0 complexes that were enriched for functional annotations. We identified 274 proteins in which we could transfer the annotation of the complex to an uncharacterized protein (**Supplemental Table 3**).

Within this set of uncharacterized proteins, we identify C7orf26 as a member of the Integrator complex (**Figure 4B**). We see strong correlation of C7orf26 with Integrator subunits in co-fractionation experiments from both HEK 293 cells and fly embryos. In addition, C7orf26 was also identified in several separate affinity purification experiments where Integrator subunits were the target bait protein (**Figure 4B**, lower panel). The consistent results from these orthogonal datasets suggest a role for C7orf26 in the Integrator complex and provides a new function for a completely unannotated protein.

We also observe another uncharacterized protein, CCDC9, as being a member of the exon-exon junction complex (EJC) (**Figure 4C**). Since the EJC is a ribonucleoprotein complex involved in RNA splicing, we searched our previously collected RNA DIF-FRAC mass spectrometry data^42^ for evidence of CCDC9 as being both a member of the EJC and also associated with RNA. The DIF-FRAC experiment identifies ribonucleoprotein complexes by comparing elution profiles of protein complexes with and without RNAseA treatment. We see in **Figure 4C** CCDC9 not only coelutes with EJC subunits but is also sensitive to RNAseA treatment suggesting it is interacting with the EJC while associated with RNA. Importantly, the DIF-FRAC experimental data was not included in the generation of hu.MAP 2.0 and therefore represents an independent assessment of CCDC9’s interaction with the EJC. Additionally, CCDC9 was identified as an RNA binding protein in high throughput screens searching for mRNA-binding proteins^43,44^ which is consistent with our observation of CCDC9 participating in a ribonucleoprotein complex.

## Conclusion

Herein, we describe the construction of the most accurate and comprehensive protein complex map to date which fills a large gap in our knowledge regarding the composition of functional complexes in the cell. We demonstrate the utility of our map by assigning functions for hundreds of completely uncharacterized proteins, providing testable hypotheses for their characterization. Additionally, we determine the prevalence of proteins that participate in multiple independent protein assemblies including ones with disparate functions suggesting moonlighting functions for the protein. Overall, our results, searchable with a simple web interface (http://humap2.proteincomplexes.org/), establish the utility of hu.MAP 2.0 for furthering our understanding of human protein functions.

### Web Resources

Website: http://humap2.proteincomplexes.org/

Code Repository: https://github.com/marcottelab/protein_complex_maps

## Acknowledgments

This work was supported by grants from the NIH (01 DK110520, R35 GM122480 to E.M.M.; R01 HL117164 and R01 HD085901 to J.B.W.; K99 HD092613 and LRP to K.D.) and the Welch Foundation (F-1515) to E.M.M.. The authors acknowledge the Texas Advanced Computing Center (TACC) at The University of Texas at Austin for providing high performance computing resources that have contributed to the research results reported within this paper. URL: http://www.tacc.utexas.edu .

## Supplemental Methods

### Mass spectrometry dataset collection

Mass spectrometry data used as input into the machine learning classifier were collected from various publications. Specifically, protein interaction features for datasets used in hu.MAP 1.0^14^ (*e.g.*, Wan *et al.*, Hein *et al.*, Huttlin *et al.*) were downloaded from http://hu1.proteincomplexes.org/static/downloads/feature_matrix.txt.gz . All HumanNet^45^ features were excluded from all model training. New datasets added for hu.MAP 2.0 were downloaded from original publications or associated dataset web resources as shown in **Table 1**.

Raw mass spectrometry data from Treiber et al.^19^ were downloaded from the Pride web resource^46^ (PXD004193) and reprocessed using the MSBlender pipeline^47^. Full details are described in Mallam et al.^42^.

HuRI dataset^23^, which was not included in training but was included for evaluation, was downloaded from http://interactome.baderlab.org/data/HuRI.tsv. HuRI protein interactions were ranked based on the number of assays the interaction was identified in.

### Weighted Matrix Model

To gain additional information on the probability that two proteins interact we generated additional features using a weighted matrix model (WMM). The WMM is based on the hypergeometric distribution and is described in Hart et al.^18^ and Drew et al.^14^ Briefly, we used the hypergeometric test (**Equation. 1**), where represents the number of experiments where both proteins A and B are identified. Variables and represent the number of experiments that independently identified protein A and protein B respectively. represents the total number of experiments.

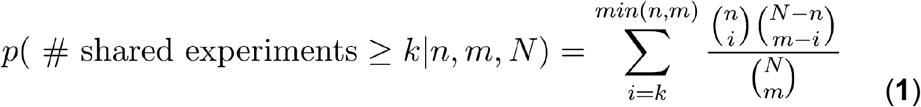

To reduce noise in the calculation, we calculated equation 1 with several cutoffs. WMM for Bioplex2.0 was calculated only considering experiments for a given protein where the protein had > 2 Bioplex2.0 Z score and > 4 Bioplex2.0 Z score. WMM based on Gupta et al. were calculated considering all experiments, > 2 average spectral counts, and > 4 average spectral counts. WMM based on Boldt et al. were calculated considering all experiments and > 4 spectral counts. WMM based on Treiber et al. were calculated considering > 2 spectral counts and > 4 spectral counts. WMM based on Youn et al. were calculated considering all experiments, > 2 spectral counts and > 4 spectral counts.

### Gold Standard Test and Training Set

To create a test and training set of literature curated protein complexes, we downloaded the complete set of Corum complexes^20^ version 2017_07_02 (http://mips.helmholtz-muenchen.de/corum/download/corum_2017_07_02.zip) and filtered out all non-human proteins. Complexes were merged to eliminate redundancy so no two complexes had > 0.6 Jaccard coefficient. Complexes were then randomly split into test and training sets. A complex was removed from the test or training sets if any pairs of proteins overlapped in the other set. Large complexes greater than 30 subunits were removed from the test and training complexes. Test and training sets were also generated for pairs of proteins for training the SVM classifier. A pair of proteins were labeled “positive” if both proteins were in the same complex. A pair was labeled “negative” if proteins were in separate complexes. All other pairs were left unlabeled. For test and training pairs, only 10% of pairs from large complexes were considered. Below is the command line used to generate the test and training sets:

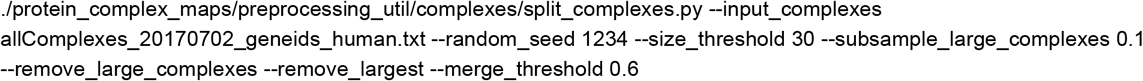

### Support Vector Machine Model Selection and Evaluation

We trained a support vector machine (SVM) classifier using Libsvm^48^ to classify pairs of proteins as co-complex protein interactions. We first generated a feature matrix using the features described above where rows are pairs of proteins and columns are features. The feature matrix was further labeled using the gold standard training set described above. We used five fold cross validation using only the training set when training to select SVM parameters C and gamma. We evaluated a range of C values (2, 8, 32, 128, 512) and gamma values (0.00048828125, 0.001953125, 0.0078125, 0.03125). As an evaluation metric, we used Area Under the Precision Recall Curve (AUPRC) averaging across the five cross validation sets. We identified C = 512 and gamma = 001953125 with the highest AUPRC. We retrained a full model using all training data using these parameters. We used this final model to predict on all pairs in the feature matrix. The final result is a list of pairs with a corresponding score generated by model.

We evaluated the final model using a precision recall framework as shown in Figure 2A. We used the scikit-learn python package^49^ to calculate precision and recall for the leave-out gold standard test protein pairs.

For comparisons between datasets as shown in Figure 2A, we generated additional models restricting the features to just those generated from the given dataset keeping the parameters C and gamma fixed. Note, the HuRI dataset was evaluated using the dataset directly as described above.

We additionally evaluated the svm confidence score for its fidelity to the test set precision value. We observed that the test set precision is consistently higher than the confidence score (**Supplemental Figure 3**). For example, a confidence score as low as 0.02 has ~0.5 precision value.

### Two-stage clustering and parameter set selection

We next used a two-stage clustering approach to identify clusters within the protein interaction network generated by the classification step described above. First, the network was thresholded based on the SVM score. We then applied the ClusterOne^24^ algorithm to identify dense regions in the thresholded network. Further, for each dense region produced by ClusterOne, we applied the MCL^25^ algorithm to identify clusters. To identify optimal parameters for the score threshold, ClusterOne parameters density and overlap as well as MCL inflation parameter, we generated clusters for various parameter combinations. Specifically, we evaluated a range of parameters: SVM score threshold (1.0, 0.99, 0.97, 0.95, 0.9, 0.8, 0.7, 0.6, 0.5, 0.4, 0.3, 0.2, 0.1, 0.05, 0.01, 0.005, 0.001, 0.0005, 0.0001, 0.00005, 0.00001), ClusterOne max overlap (0.6, 0.7, 0.8), density (0.1, 0.2, 0.3, 0.35, 0.4) and MCL inflation (1.2, 2, 3, 4, 5, 7, 9, 11, 15). We also compared using an unweighted graph as input into ClusterOne versus a weighted graph and observed the unweighted graph had superior performance. A weighted graph was used for the MCL stage clustering. We additionally applied a post-clustering filter that removed nodes from a cluster that lacked edges that scored greater than the SVM score threshold.

To evaluate the clusterings as shown in Figure 2B and 2C, we used the *k*-cliques method, specifically weighted recall (R_weighted) and weighted precision (P_weighted), which we described previously^14^. Briefly, the *k*-cliques method globally compares a set of clusters to a set of gold standard complexes by comparing cliques derived from the clusters to cliques derived from gold standard complexes. This comparison is done for all clique sizes from size 2 (ie. pairs) to size n (ie. the size of the largest complex or cluster). A precision and recall value are calculated for all clique sizes. A weighted average is then calculated for both precision and recall across all clique sizes, weighted by the number of clusters with size >= to the clique size.

We evaluated all clusterings using the *k*-cliques method, comparing to the training set of gold standard complexes. We selected five clusterings that optimize the tradeoff between precision and recall as shown in **Figure 2B**. These five clusterings were then combined into a union set. **Table 2** shows the clustering parameters used for the selected clusterings.

We finally evaluated the individual selected clusterings as well as the union of the selected clusterings using the *k*-clique method by comparing to the leave-out set of gold standard test complexes (Figure 2C). In addition, we compare to previously published complex maps from Wan et al.^10^, Bioplex 1.0^7^, Bioplex 2.0^8^, as well as our original hu.MAP 1.0^14^.

### Identification of Promiscuous Proteins

To identify proteins which participate in multiple complexes, we first determined a set of complexes with limited overlap. To determine the degree with which one complex overlaps another we developed a ‘subcomplex index’ (**Equation 2**) defined as:

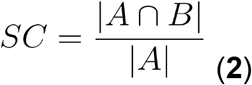

where *A* and *B* are complexes (ie. sets of proteins). The ‘subcomplex index’ is related to the Jaccard index but is normalized by the size of a single complex rather than the size of the union of both complexes as is done in the Jaccard index. We then calculated the subcomplex index for every complex compared to all other complexes. We then generated a set of complexes with limited overlap by selecting complexes that had SC < 0.5 to all other complexes. We then identified all proteins that participated in multiple complexes from this reduced set of complexes.

### Calculation of protein age enrichment for promiscuous proteins

Protein ages were mapped using 'modeAge' in the main_HUMAN.csv file from Liebeskind et al.^50^. Z-scores for each age group were determined by comparing the number of promiscuous proteins to a background distribution. The background distribution was calculated by counting the number of randomly sampled non-promiscuous proteins (i.e. proteins that participate in only 1 complex) in each age group.

### Annotation Enrichment

Annotation enrichment was calculated for GO, Reactome, CORUM, KEGG and Human Phenotype Ontology (HP) terms using gProfiler^27^ for each individual complex. All proteins observed in the 15,000 mass spectrometry experiments were used as the background set. Annotations inferred by electronic transfer were ignored.

To evaluate annotation enrichment for all complexes, we first generated a set of shuffled complexes where protein ids were reassigned to new cluster ids. This has the effect of keeping both the number of clusters and the size distribution of clusters the same as the final set of hu.MAP 2.0 complexes. Annotation enrichment for the shuffled set of clusters was done as described above. Using this background annotation enrichment from all categories, we calculated a 0.05 false discovery rate threshold.

## Supplemental Figures

**Supplemental Figure 1.**
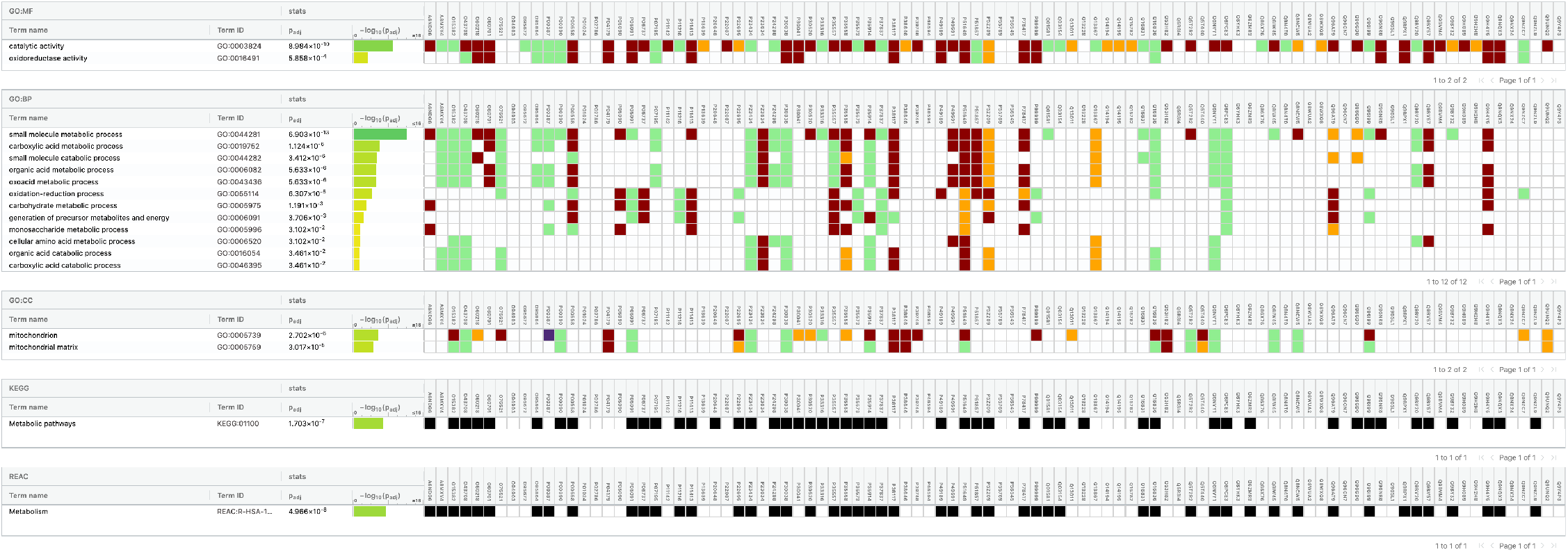
Annotation enrichment of older promiscuous proteins. gProfiler output shows older promiscuous proteins are enriched for metabolic processing annotations.

**Supplemental Figure 2.**
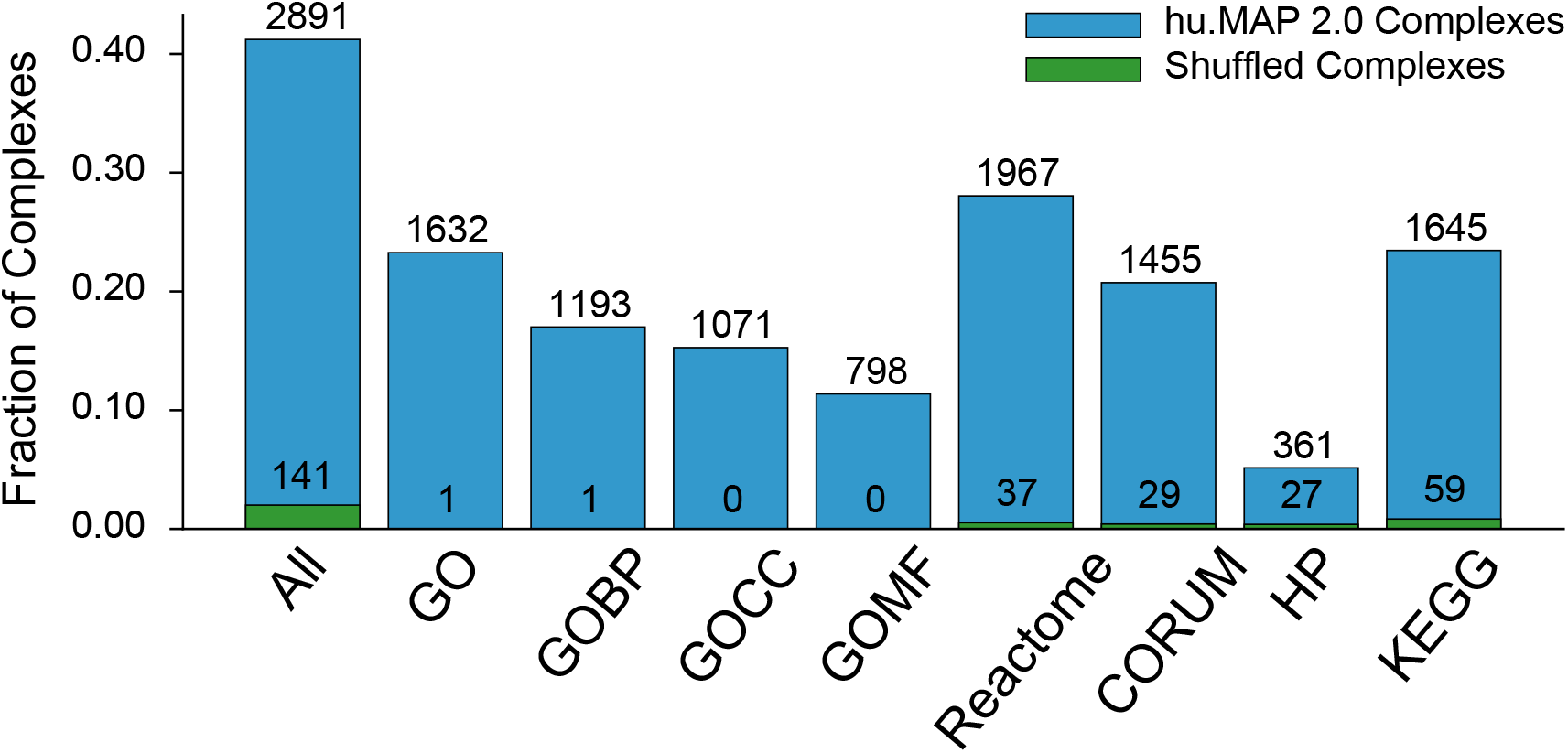
hu.MAP 2.0 complexes are functionally enriched. The bar chart shows the number of identified complexes that are enriched with at least one annotation from GO, Reactome, CORUM, KEGG or Human Phenotype Ontology (HP) at an FDR threshold of 0.05.

**Supplemental Figure 3.**
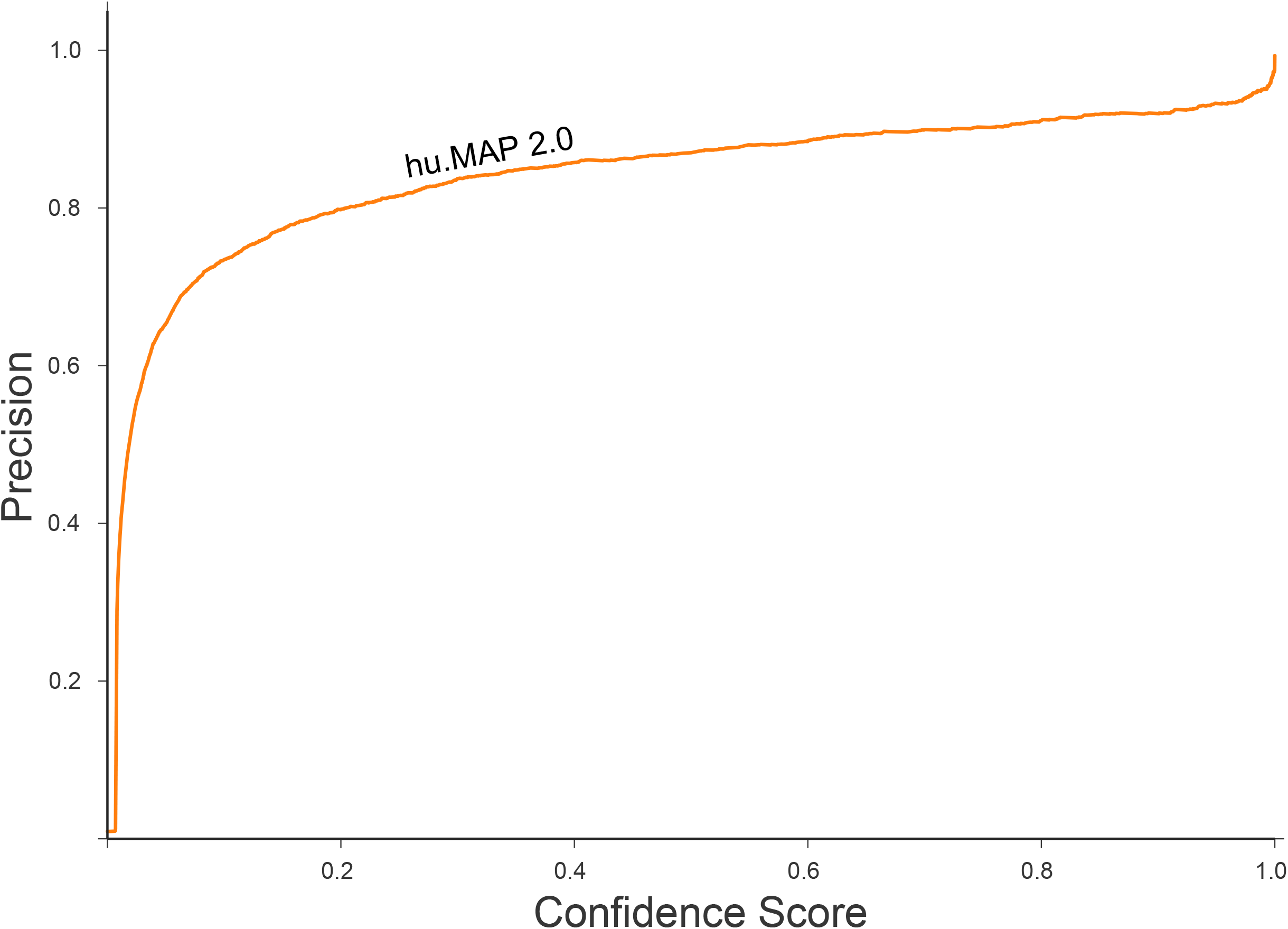
SVM confidence score versus test set precision. The line plot shows the relationship between the SVM confidence score and the empirical precision value calculated from the test set of protein interactions. The relationship shows the precision value is consistently higher than the confidence score.

## Supplemental Tables

**Supplemental Table 1. Table of hu.MAP 2.0 protein complexes**

**Supplemental Table 2. Table of promiscuous proteins**

**Supplemental Table 3. Table of identified annotations for uncharacterized proteins**

